## Supplementary Table 1 for "A systematic examination of preprint platforms for use in the medical and biomedical sciences setting"

**Supplementary Table 1: Reasons for excluded preprint platforms**

| **Name of Site** | **Reason for Exclusion** |
| --- | --- |
| Chemweb | Inactive |
| Centre for Health Economics and Policy Analysis (CHEPA) | Inactive |
| ClinMed NetPrints | Inactive |
| Cogprints | Inactive |
| CSTC | Inactive |
| K-Theory Preprint Archives | Inactive |
| Mathematics Preprint Search System (MPRESS) | Inactive |
| National Advisory Committee for Aeronautics (NACA) | Inactive |
| Nature Precedings | Inactive |
| The Winnower | Inactive |
| Geometry Center's Preprints | Inactive |
| World Health Organisation Zika Open repository | Inactive |
| Instant Math Preprints (Yale Mathematics Preprint Bulletin Board) | Inactive |
| Organisation Européenne pour la Recherche Nucléaire document platform | Repository |
| Hyper Articles en Ligne (HAL) | Repository |
| ResearchGate | Repository |
| White Rose Consortium e-Prints Repository | Repository |
| Zenodo | Repository |
| Organisation Européenne pour la Recherche Nucléaire Print | Offline |
| Fermilab | Offline |
| Physics Information Exchange (PIE) | Offline |
| Radio Astronomy Preprints – RAPsheet | Offline |
| Space Telescope Preprints – STEPsheet | Offline |
| Advance: a SAGE preprints community | Scope: Humanities and Social Sciences |
| BodoArXiv | Scope: Medieval Studies |
| Cryptology ePrint Archive | Scope: Cryptology |
| CORE repository | Scope: Humanities |
| EarthArXiv | Scope: Earth Sciences |
| EconStor | Scope: Economics and Business Studies |
| ECSarXiv | Scope: Electrochemistry and solid state science |
| engrXiv | Scope: Engineering |
| E-LIS | Scope: Library and Information Science |
| Electronic Colloquium on Computational Complexity | Scope: Computer Science |
| Institute for Fiscal Studies (IFS) Working Papers | Scope: Economics |
| LawArxiv | Scope: Law |
| LIS Scholarship Archive | Scope: Library and Information Science |
| LingBuzz | Scope: Linguistics |
| MediArXiv | Scope: Media, Film and Communication Studies |
| Mathematical Physics Preprint Archive (mp_arc) | Scope: Mathematical Physics and Related Areas |
| National Bureau of Economic Research (NBER) Working Papers | Scope: Economics |
| Networked Computer Science Technical Reference Library (NCSTRL) | Scope: Computer Science |
| Philsci Archive | Scope: Philosophy of Science |
| Munich Personal RePEc Archive (MPRA) | Scope: Economics |
| Social Science Open Access Repository (SSOAR) | Scope: Social Sciences |
| Stanford Physics Information Retrieval System (SPIRES) | Scope: Physics |
| WorldBank's Policy Research Working Paper Series (PRWPs) | Scope: Economics |
